# Mixture Models Unveil the Origin of the Enigmatic Satyrine Butterfly Genera *Calisto* and *Llorenteana* (Lepidoptera, Nymphalidae, Satyrinae)

**DOI:** 10.64898/2026.02.01.703087

**Authors:** Rayner Núñez, Anja Bodenheim, Yosiel Alvarez, Niklas Wahlberg, Marianne Espeland

**Affiliations:** Leibniz Institute for the Analysis of Biodiversity Change –Museum Koenig, Bonn, 53113, Germany; Laboratório de Ecologia e Sistemática de Borboletas, Instituto de Biologia, Universidade Estadual de Campinas, Campinas, 13083-865, Brazil; Department of Biology, Lund University, Lund, 223 62, Sweden

**Keywords:** biogeography, divergence times, gene flow, incomplete lineage sorting, phylogenetic relationships, Satyrini, systematics

## Abstract

We provide the first comprehensive analysis of the origin of two enigmatic Satyrinae genera of uncertain affinities. *Calisto*, the only Satyrinae genus from the West Indies and endemic to these islands, has resisted numerous attempts at phylogenetic placement, regardless of the data type or methods used. *Llorenteana*, a monotypic genus from northwestern Mexico, has never been included in a molecular phylogenetic study, and past authors have placed it in five different genera and subtribes. We used mostly published genomic data, but also newly sequenced whole genome data from museum specimens and old DNA extracts, extracted BUSCO genes and prepared several datasets. These datasets differed in the degree of heterogeneity and saturation, the number of nucleotide positions used (all positions or only the first two), and were analyzed as nucleotides or as amino acids. We employed several methods for phylogenetic reconstruction using both partitioned and mixture models, as well as ASTRAL, and we inferred divergence times and ancestral areas of origin for *Calisto* and *Llorenteana*. The phylogenetic placement of *Calisto* varied among datasets when we used partitioned models and ASTRAL; however, most datasets resulted in the same relationships under mixture models. Our results suggest that *Calisto* is part of a clade of Old World origin that colonized the New World from north to south, thus sharing ancestry with Nearctic taxa. *Llorenteana* constitutes one of the earliest splits within the Euptychiina, a subtribe of Neotropical origin, but descending together with the Pronophilina from Nearctic ancestors. We propose the recognition of Erebiina **stat. rev.** as the only subtribe comprising the former Calistina **syn. nov.**, Callerebiina **syn. nov**, Maniolina **syn. nov.**, and Ypthimina **syn. nov.**

## Introduction

The subfamily Satyrinae, with more than 2800 known species (Peña and Wahlberg 2025), comprises approximately 40% of the family Nymphalidae, representing one of the largest radiations among butterflies. Although this group has a worldwide distribution, the Neotropical region harbors approximately half of this diversity (Lamas 2004). Until the early 2000s, there was widespread confusion regarding the subfamily’s status and the relationships among its clades (Peña et al. 2006). The advent of molecular phylogenetic studies has shed light on the relationships within Satyrinae. The use of Sanger sequencing, and more recently of high-throughput sequencing, has helped to clarify the status and the relationships among most larger clades within the subfamily (Murray and Prowell 2005; Peña et al. 2006; 2010; 2011; Peña and Wahlberg 2008; Wahlberg et al. 2009; Kodandaramaiah et al. 2010; Matos-Maraví et al. 2014; Aduse-Poku et al. 2015; Espeland et al. 2018; 2019; 2023; Chazot et al. 2019; Kawahara et al. 2023). It is now acknowledged that the Satyrinae comprise nine monophyletic tribes: Brassolini, Morphini, Amathusiini, Zetherini, Dirini, Elymniini, Haeterini, Melanitini and Satyrini, the latter including approximately 80% of the species.

The relationships within Satyrini, however, have remained problematic; the subtribe structure is still unclear, and several genera remain to be placed (Viloria and Luis-Martínez 2019; Espeland et al. 2023; Kawahara et al. 2023). Peña et al. (2010; 2011) suggested long-branch attraction as a possible cause of these inconsistencies. They and other authors (*e.g.* Espeland et al. 2018; 2023) also noted that the rapid radiation undergone by the Satyrini, evidenced by the short branches leading to many major clades, has likely obscured their relationships. Some of these works (*e.g.* Espeland et al. 2018; 2019, Kawahara et al. (2023) performed coalescent analyses and removed loci with high guanine-cytosine (GC) content to account for incomplete lineage sorting and compositional bias. Rate heterogeneity among lineages is another factor that might affect the phylogenetic inference, and which has so far not received much attention in this system. The traditionally used partitioned models assume that substitution rates are constant at each site across all lineages, and it has been shown that heterotachously-evolved sequences can lead to model misspecification (Kolaczkowski and Thornton 2004; Som 2015).

The genus *Calisto* Hübner is one of the most enigmatic genera within the Satyrinae. It is the only representative of the subfamily in the Greater Antilles. The islands are relatively old, with estimates of emerged land in the region dating back to the late Eocene, approximately 35 Ma (Iturralde-Vinent, 2006). Since then, the archipelago has undergone numerous transformations driven by tectonic movements and changes in sea level and climate, shaping the colonization-extinction dynamics of the islands (Ricklefs and Bermingham 2007). The current biota of the islands is characterized by radiations and high endemism in groups such as land snails (Proios et al. 2021), *Anolis* lizards (Muñoz et al. 2023), *Eleutherodactylus* frogs (Jiménez-Ortega et al. 2023), and vascular plants (Nieto-Blázquez et al. 2017). Owing to these and several other radiations, the region has been included among the world’s biodiversity hotspots (Myers et al. 2000; Mittermeier et al. 2005).

At the same time, the processes mentioned above, along with the islands’ isolation, size, height, and resource availability, among other factors, have resulted in the depauperation or complete absence of other lineages that are highly diverse on the continent. Well-known examples include freshwater fishes (Massip-Veloso et al. 2024) and terrestrial mammals (Dávalos 2004). However, many other cases of missing or depauperate groups occur on the islands, including invertebrates such as insects, butterflies among them. For instance, the Hesperiidae subfamily Pyrrhopyginae is absent, the Nymphalidae tribe Ithomiini and the family Riodinidae are represented each by a single extant genus, and the species-rich genera *Parides* Hübner (Papilionidae) and *Heliconius* Kluk (Nymphalidae) have each only one species in the islands (Smith et al. 1994). The genus *Calisto* exemplifies both of these contrasting patterns. On the one hand, *Calisto* is the sole member of the Satyrinae in the archipelago; on the other, it represents the largest known Lepidoptera radiation in the islands, with more than 50 described species (Núñez et al. 2019).

In the phylogenetic studies cited above, those including *Calisto* always recovered the genus within the tribe Satyrini, regardless of the data type, sampling size, or inference method employed. However, the closest affinities of *Calisto* have remained elusive. In many cases, the relationships lacked support, with *Calisto* found at different places in the phylogeny (Fig. S1).

*Llorenteana* Viloria and Luis-Martínez is another Satyrinae pending phylogenetic assignment. The genus is monotypic and its single species has previously been included in five genera in different subtribes (Viloria and Luis-Martínez 2019). *Llorenteana pellonia* (Godman) inhabits a small area of northwestern Mexico, within the Neotropical region but near its boundary with the Nearctic. This taxon has never been included in a molecular phylogeny, and according to Espeland et al. (2023) likely belongs in the largely neotropical subtribe Euptychiina, but Viloria and Luis-Martínez (2019), based on morphology, suggested placement in the old-world subtribe Ypthimina.

Here, we aimed to infer the phylogenetic position of *Calisto* and *Llorenteana* by analyzing genomic data using several tree inference methods. Given the different placements recovered for *Calisto* in previous studies, we also accounted for possible sources of phylogenetic incongruence such as rate heterogeneity, saturation, compositional bias, incomplete lineage sorting, and gene flow (Scornavacca et al. 2020; Kapli et al. 2021; Fleming et al. 2023). We prepared several datasets containing between 391 and 2580 BUSCO genes of representatives of all Satyrini subtribes and outgroups. We analyzed the genes for compositional bias and saturation and repeated the tree inference process after removing the affected loci. We also evaluated the extent to which incomplete lineage sorting and ancestral gene flow affect our data and the inferred trees. In addition to the traditional partitioned models, we reconstructed the relationships using nucleotide mixture models (Ren et al. 2025), applied for the first time in this group. We inferred the biogeographical history of *Calisto* and *Euptychia*. We used a recently published dataset (Kawahara et al. 2023), expanded to include additional species of both genera, representing the largest genomic sampling of Satyrini to date.

## Material and Methods

### Sampling and Datasets Preparation

Our main dataset contained 69 representatives of all Satyrini subtribes and three outgroups (Table S1). Of these, we downloaded 31 genome assemblies and 37 raw data samples from NCBI. Of the genome assemblies, 22 of them generated as part of the Darwin Tree of Life project (The Darwin Tree of Life Project Consortium 2022) and Project Psyche (Wright et al. 2025) (Table S1).

We sequenced medium-coverage (aiming for 15 X) genomes for three *Calisto* species and a putative member of the Euptychiina never sampled before, the genus *Llorenteana*. We extracted the latter from a museum specimen collected in 1984, deposited at the McGuire Center for Lepidoptera and Biodiversity, using the QIAGEN DNAeasy Blood and Tissue Kit. We measured DNA concentration and fragment length for this sample, and three 12-year-old Calisto DNA extracts from Matos-Maraví et al. (2014), using Quantus (Promega) and Fragment Analyzer (Agilent), respectively. We used the Library preparation kit NEBNext Ultra II DNA Library Prep kit (New England Biolabs) and sequenced them using Illumina 150 bp paired end reads (Macrogen). We assembled the resulting raw as well as the SRA files downloaded from GenBank using SPAdes 3.15.5 with default parameters (Prjibelski et al. 2020).

We extracted BUSCO genes using the buscophy workflow (https://gitlab.leibniz-lib.de/smartin/buscophy) with all newly produced assemblies, including those obtained from NCBI SRA data, and reference genome assemblies downloaded from NCBI. This workflow searches each assembly for all single copy BUSCO genes, complete or fragmented, present in a reference database, OrthoDB v10 in our case (Kriventseva et al. 2019). Subsequently, the workflow aligns the extracted amino acid data using MAFFT-linsi (Katoh et al. 2002) and converts these alignments together with their corresponding nucleotide sequences to codon-based DNA alignments using pal2nal (Suyama et al. 2006). Only loci with a minimum number of samples are included, 50% of the samples in our case. We used OliInSeq v0.9.6 (parameters -M -w 8 -f 2 -R) to remove outliers (https://github.com/cmayer/OliInSeq), and trimAl (Capella-Gutierrez et al. 2009) to remove any column with more than 50% missing data. Each nucleotide alignment was visually inspected in Aliview 1.28 (Larsson 2014) and finally loci shorter than 200 bp were removed. We used PhylteR (Comte et al., 2023) to detect any remaining potential outliers, entire genes, or specific samples within them, applying different values for the ‘k’ parameter (3, 4, and 6) that controls the detection strength. We only analyzed in depth the results of the most strict run (k=3) that highlighted no gene outliers and 121 outlier sequences in 35 genes. These results implied a data loss of 0.1%; however, after manually checking all outlier gene trees and their respective fasta files, we removed only 27 sequences from 14 genes. Most highlighted sequences correspond to *Calisto*, which usually sits at the end of a long branch, likely the reason they were highlighted by PhylteR.

We prepared several datasets and obtained summary statistics using AMAS (Borowiec 2016): nucleotides (NT123), the first two codon positions (NT12), and amino acids (AA). For the latter, we translated the cleaned and filtered nucleotide alignments to amino acids using FASconCAT-G (Kück and Longo 2014). Other datasets were prepared according to different levels of saturation and compositional bias (Table S2): NT123 low saturation (NT123 LOWSAT), and others renamed after the thresholds used to remove genes with high compositional heterogeneity: NT123 0.025/0.004, NT12 0.001/0.0015, and AA 0.0025/0.004 (Table S2). We prepared another dataset, NT123 KW, containing 46 Satyrinae, non-Satyrini members, and all 233 Satyrini from Kawahara et al. (2023), plus one additional *Gyrocheilus* and two more species of *Euptychia* and *Calisto*, and one sample of *Llorenteana pellonia* (Godman) to posteriorly perform ancestral range reconstruction analyses.

### Compositional Bias and Saturation

We used BaCoCa v 1.1 (Kück and Struck 2014) to analyze our nucleotide and amino acid sequence alignments and calculate their compositional heterogeneity and saturation. First, the program estimates relative composition frequency variability (RCFV); however, we calculated an improved version called normalized RCFV (nRCFV), available in RCFV_reader (Fleming and Struck 2023), which avoids the biases of the original metric: sequence length, number of taxa, and number of possible character states within the dataset. For the second metric, BaCoCa calculates the c value (convergence), which is the ratio of the standard deviation of all transition to transversion ratios and the standard deviation of the uncorrected genetic p distances. The standard deviation of the transition to transversion ratio decreases when saturation increases, while the ratio of uncorrected genetic p distances also increases. Lower c values indicate a higher degree of saturation. We performed principal component analyses (PCA) in PAST 4.02 (Hammer et al., 2001) to explore the base frequency variation of all nucleotide positions and amino acids for *Calisto* and its closest relatives.

### Phylogenetic Inference

We applied a maximum likelihood approach using IQ-TREE 2.2.6 (Minh et al. 2020) to our concatenated datasets. We used ModelFinder (Kalyaanamoorthy et al. 2017) to select the best partition model per gene for the nucleotide (-m MFP) and amino acid (–msub nuclear) datasets. We calculated branch support with 1000 ultrafast bootstraps (UFBoot, - B 1000) (Hoang et al. 2018) and used the nearest-neighbor interchange additional step (-bnni) to further optimize UFBoot trees and avoid overestimating branch supports due to severe model violations. For each dataset/analysis five tree searches were performed (--runs 5), and the tree with the highest likelihood selected as the best tree.

We reanalyzed all nucleotide datasets in IQ-TREE using mixture models (Pagel and Meade 2005) inferred with MixtureFinder (Ren et al. 2025). These models minimize assumptions (e.g., heterotachy only occurs between partitions, not within each partition) and thus are less prone to model misspecification. MixtureFinder incrementally adds one class after another, each with its own substitution model (inferred by ModelFinder) and rate heterogeneity across sites, until finding the optimal number of classes for the dataset meaning that adding a new class will not improve the fit. However, this approach implies that mixture model search becomes more computationally intensive with each new class addition. The large size of our datasets, compared to the empirical and simulated ones employed to assess MixtureFinder performance by Ren et al. (2025), prevented the completion of our model search runs. The analyses crashed at the fifth, sixth or seventh class before finding the optimal number of classes and their models. Thus, we used the 4-class model inferred for each dataset, as the results by Ren et al. (2025) show that mixture models outperform partitioned models starting from the second class. We used the default Bayesian Information Criterion since tests have shown that mixture model selection is relatively insensitive to the criterion applied (Ren et al. 2025). We also applied another mixture model, GHOST (Crotty et al. 2020), to all nucleotide datasets. This model runs with the parameters linked (common) or unlinked (separate) for each class. We used a 4-class GHOST model to infer all parameters (branch lengths, substitution rates, inferred base frequencies) independently for four classes for each dataset, GTR+FO*H4. This model performs better than traditional partitioned models if heterotachy (the change in evolutionary rates across lineages) affects the dataset. For the amino acid datasets, we applied the mixture models LG, LG4X, and PMSF (Le and Gascuel 2008; Le et al. 2012; Wang et al. 2018).

We also inferred coalescent species trees in ASTRAL-IV v1.23.4.6 (Zhang et al. 2025), which integrates CASTLES-II (Tabatabaee et al. 2023) and computes terminal and internal branch lengths in substitution-per-site units. For these reconstructions, we used gene trees inferred in IQ-TREE using one partition model per locus and substitution models selected using ModelFinder as above.

We calculated gene concordance factors (gCF) and site concordance factors (sCF) using IQ-TREE 2 (Minh et al. 2020; Mo et al. 2023).

### Incomplete Lineage Sorting and Ancestral Gene Flow

We investigated whether incomplete lineage sorting (ILS) and ancestral gene flow (GF) affect our phylogenetic inference process by using aphid (Galtier 2024). This program analyzes the topologies and branch lengths of rooted gene trees of a given species triplet, *e.g.* ((A, B) C), using a maximum likelihood framework. In our case, each triplet included one *Calisto* representative, one Pronophilina or Euptychiina, and one member of the Erebiina herein including also Callerebiina, *Gyrocheilus*, Maniolina, and Ypthimina all of which grouped into the same clade (see Results). We performed the analysis three times setting the most likely topology according to the phylogenetic inference results as the main topology plus two other alternative topologies. We generated a script to obtain all possible triplets of the three groups present in our main dataset, NT123, totaling 1008. After discarding trees lacking *Morpho helenor* (used for rooting and a requisite) and all non-monophyletic triplets, the program analyzed the remaining triplets with the default parameters. Aphid classifies the gene trees into five categories: 1) posterior no event, the gene tree matches the main topology and branching times, 2) posterior no conflict ILS, same topology, but branching times are higher than in the main topology, 3) posterior no conflict GF, same topology, but branching times are lower than in the main topology, 4) posterior conflict ILS (ILSc), topology differs from the main topology and branching times are higher, 5) posterior conflict GF (GFc), topology differs from the main topology and branching times are lower. We only analyzed triplets containing 50 or more rooted gene trees with a monophyletic triplet.

### Divergence Times and Ancestral Range Estimation

We estimated the divergence times for the Satyrini using trees inferred with the main, NT123 (partitioned and GHOST model), and the NT123 KW datasets with the GHOST and a 4-class mixture model. Since no fossils have been reliably assigned to a Satyrinae clade, we used secondary calibration points from a published dated butterfly phylogeny. We used the ages and the 95% confidence intervals estimated by Espeland et al. (2018) to set three calibration points for the NT123 dataset and five for the NT123 KW dataset, respectively (File S1). Despite their larger sampling, we did not use the age estimates by Kawahara et al. (2023) obtained with TreePL since apparently this maximum likelihood method leads to older divergence times (Costa et al. 2022), which can lead to conflict with the fossil record (Schachat et al. 2023). We used RelTime-ML (Tamura et al. 2012) in MEGA12 (Kumar et al. 2024) to estimate the divergence times using a GTR+I+G model and normal distribution priors. This maximum likelihood method has proven to be more computational efficient and gives results concurring or closer to those of Bayesian methods (Mello et al. 2021; Costa et al. 2022).

We performed ancestral range reconstruction analyses for *Calisto* and the entire tribe Satyrini using BioGeoBEARS (Matzke, 2013), which implements the DEC, DIVALIKE and BAYAREALIKE models within a maximum likelihood framework. We subdivided the Satyrini geographical range into seven areas: A) Palearctic, B) Nearctic, C) Neotropics excluding the West Indies, D) Afrotropics, E) the Indomalayan region, F) Australasia, and G) the West Indies, including the Greater Antilles, the Bahamas and Anegada Island (the entire range of *Calisto*) (Supplementary file 2). For the analyses we used the time calibrate tree obtained in RelTime with the NT123 KW dataset. We performed unconstrained and constrained analyses setting dispersal probabilities for the latter according to the position of each area respect the others (File S2), as well as a two time slices, before and after 35 Ma, which, according to Iturralde and MacPhee (1999), marks the presence of permanent land at the area occupied by the archipelago. We used the Akaike information criterion corrected for sample size (AICc) to compare results of the different models.

## Results

### Datasets, Compositional Bias and Saturation

We prepared nine datasets containing between 391 and 2,580 loci and between 161,166 and 2,587,844 base pairs, respectively (Table S2). The percentage of informative sites and missing data ranged from 12 to 47% and from 11 to 37%, respectively. Some of the datasets obtained after filtering out genes with high nRCFV values had considerable lower amounts of informative sites, e.g. NT12 0.001, while others kept values relatively close to the parent dataset, *e.g.* NT123 0.004/0.0025 (Table S2, File S3).

The saturation c-convergence values differed markedly between two groups of genes that exhibited similar nRCFV ranges (Fig. S2). We created the NT123 LOWSAT dataset with the group of genes with the lowest saturation (higher c values). File S3 contains the summarized output of the BaCoCa analyses for each dataset, for the entire alignment and per gene, and for each taxon. The PCA analysis of the base frequencies for all nucleotide positions and amino acids revealed they are highly variable within Euptychiina and within *Calisto* and its sister clade compared to the Satyrina and Pronophilina clades (Fig. S3, File S3). As expected, the third codon position contributed the most to the observed variation. However, the analysis including only the first two codon positions also revealed a high variability within the same clades (Fig. S3).

### Phylogenetic Inference

The topologies of our inferred trees vary depending on the dataset and the type of model employed. Regarding the subtribes not closely related to *Calisto*, we obtained rather constant topologies, except for the subtribe Ragadina (*Acrophtalmia* and *Ragadia*), sister to all other subtribes in some trees or sister to the subtribes Mycalesina and Lethina in others.

In most of our trees, *Calisto* is part of a clade that includes the subtribes, Callerebiina, Erebiina, *Gyrocheilus*, Maniolina, and Ypthimina (Figs. 1, 2, File S4). We obtained weak support for some of the relationships among these clades in most trees inferred with partitioned models with the Ypthimina being paraphyletic in several of them. Only the NT123 dataset and its versions with reduced compositional bias, 0.004/0.0025, yielded fully resolved relationships between *Calisto* and its closest relatives (Fig. 2, File S4).

**Figure 1.**
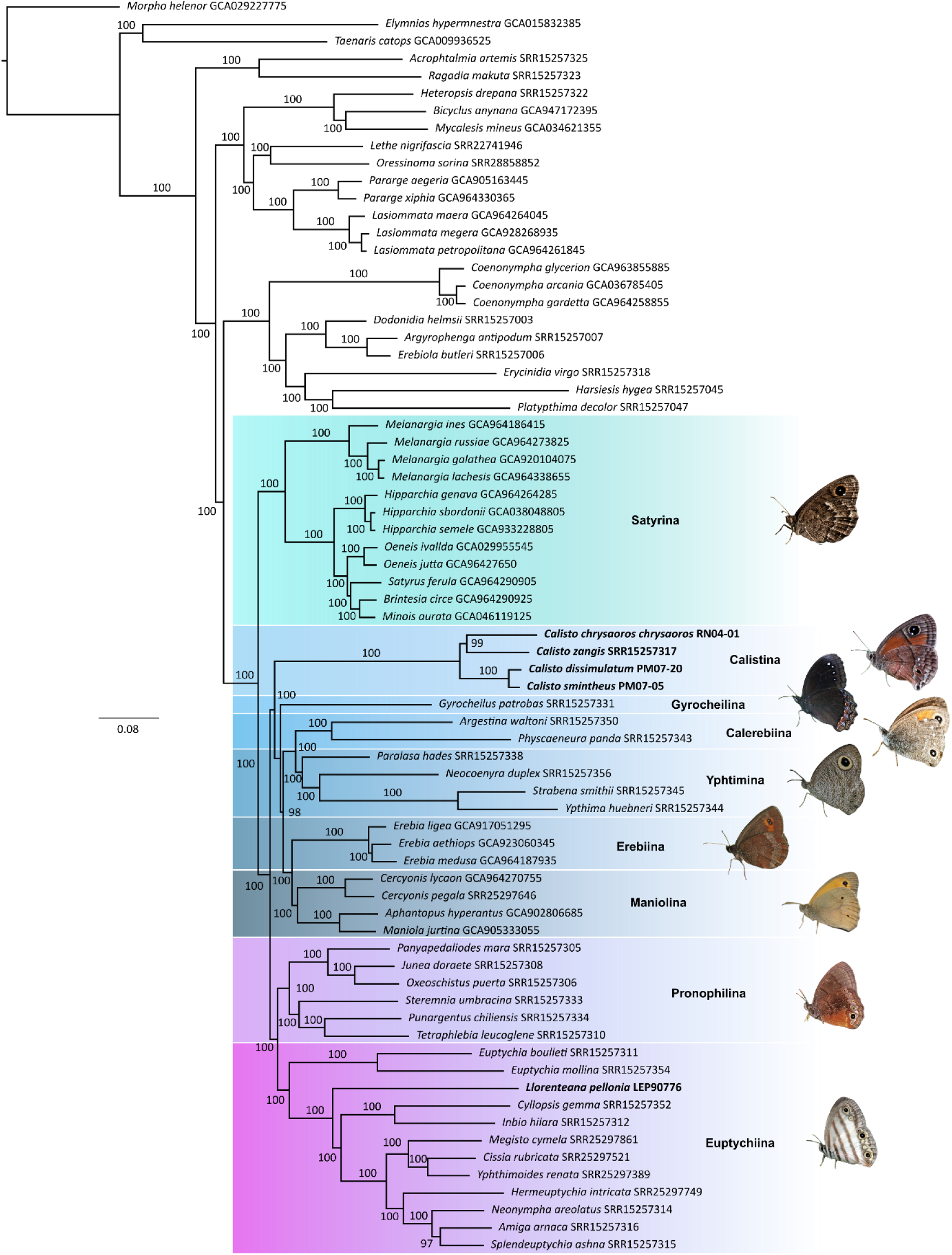
Maximum likelihood best tree inferred in IQ-TREE with the NT123 dataset and the GHOST model showing the relationships of the genera *Calisto* and *Llorenteana* within the tribe Satyrini. Numbers at the branches represent IQTREE ultrafast bootstrap values. Color boxes represent the subtribes closely related to *Calisto* and *Llorenteana*. Both genera and related subtribe names shown in bold. Butterflies images credits: A. R. Perez-Asso, Maarten Sluijter, Janet Graham, Oleg Kosterin, Zhong Yankui, Charles J. Sharp, Paul Prappas, Anna N. Chapman, and desertnaturalist.

**Figure 2.**
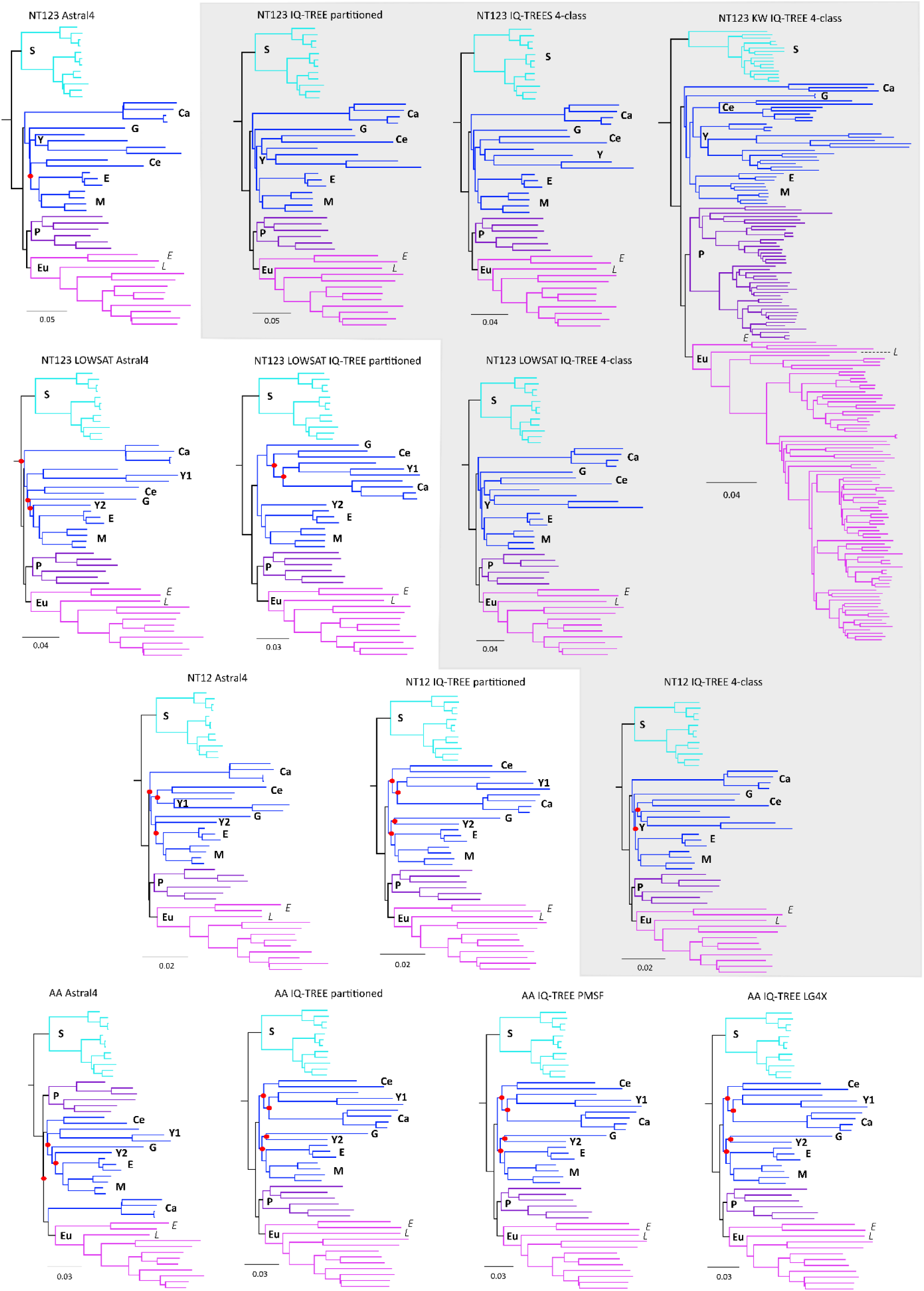
Relationships of *Calisto* and *Llorenteana* and their closest relatives within the tribe Satyrini inferred using different reconstruction methods, datasets and models. The gray polygon highlights trees with the same topology. Red dots mark weakly supported branches for *Calisto* and allied taxa. Colors identify major clades; a single color represents Calistina, Callerebiina, Erebiina, Gyrocheilina, Maniolina and Yphtimina. Abbreviations: Calistina (Ca), Callerebiina (Ce), Erebiina (E), *Euptychia* (E), Euptychiina (Eu), *Llorenteana* (L), Gyrocheilina (G), Maniolina (M), Pronophilina (P), Satyrina (S), and Yphtimina (Y, numbers represent paraphyletic splits). These and all other trees are available in File S4.

The Astral-IV trees inferred using the NT12 0.0015 and AA datasets and partitioned models place *Calisto* as sister to Euptychiina with strong support while the relationships among the other clades lack support. Other strongly supported unlikely placements included *Paralasa* (Ypthimina) as sister to the Erebiina-Maniolina in the N123 LOWSAT IQ-TREE tree, and *Gyrocheilus* as sister to *Paralasa* plus Erebiina-Maniolina in the NT12 Astral-IV tree.

The relationships inferred with all nucleotide datasets using a 4-class mixture model converged on the same topology. Pronophilina and Euptychiina are sister groups and this clade is sister to another clade where *Calisto* is sister to the remaining groups (Figs. 1, 2, File S4). In this topology, *Calisto* is sister to *Gyrocheilus* and a clade that comprises Erebiina-Maniolina as sister to Callerebiina-Ypthimina. Furthermore, these results concur with the topology inferred using the GHOST model across most datasets, also with strong support. Only the least informative datasets, NT12 0.001/0.0015, produced trees with several weakly supported paraphyletic placements. The three protein mixture models applied to the AA dataset found strong support for the sister relationship between Pronophilina and Euptychiina, but weak support for paraphyletic relationships among the remaining clades (File S4). None of the trees inferred using mixture models produced paraphyletic strongly supported relationships for *Calisto* and its closely related taxa.

The relationships between *Calisto* and its closest allies, as inferred with the NT123 KW dataset and a 4-class mixture model, match the aforementioned also with strong support (Fig. 2, File S4). However, the partitioned models found *Calisto* as sister to *Gyrocheilus* with strong branch support (File S4).

In all trees, the placement of the genus *Llorenteana* is the same, as a member of the subtribe Euptychiina (Figs. 1, 2, File S4). Its sole representative, *L. pellonia*, constitutes the first branch after the genus *Euptychia* (the earliest split within the subtribe), and is sister to the rest of the Euptychiina.

### Trees Comparison and Concordance Factors

Comparing the branch lengths and support of the inferred trees provides insight into the performance of the different models. For several datasets, the four-class mixture models, especially the GHOST model, produced longer trees with longer inner branch lengths (Fig. 3). In several cases, these trees have higher average branch support when analyzing either the entire tree or only *Calisto* and its closely related clades. The number of weakly supported branches within the latter group decreased in several trees inferred with the 4-class and GHOST mixture models (Fig. 3). The Astral-IV trees have the shortest inner branch lengths of all the trees, primarily at the splits among groups closely related to *Calisto*.

**Figure 3.**
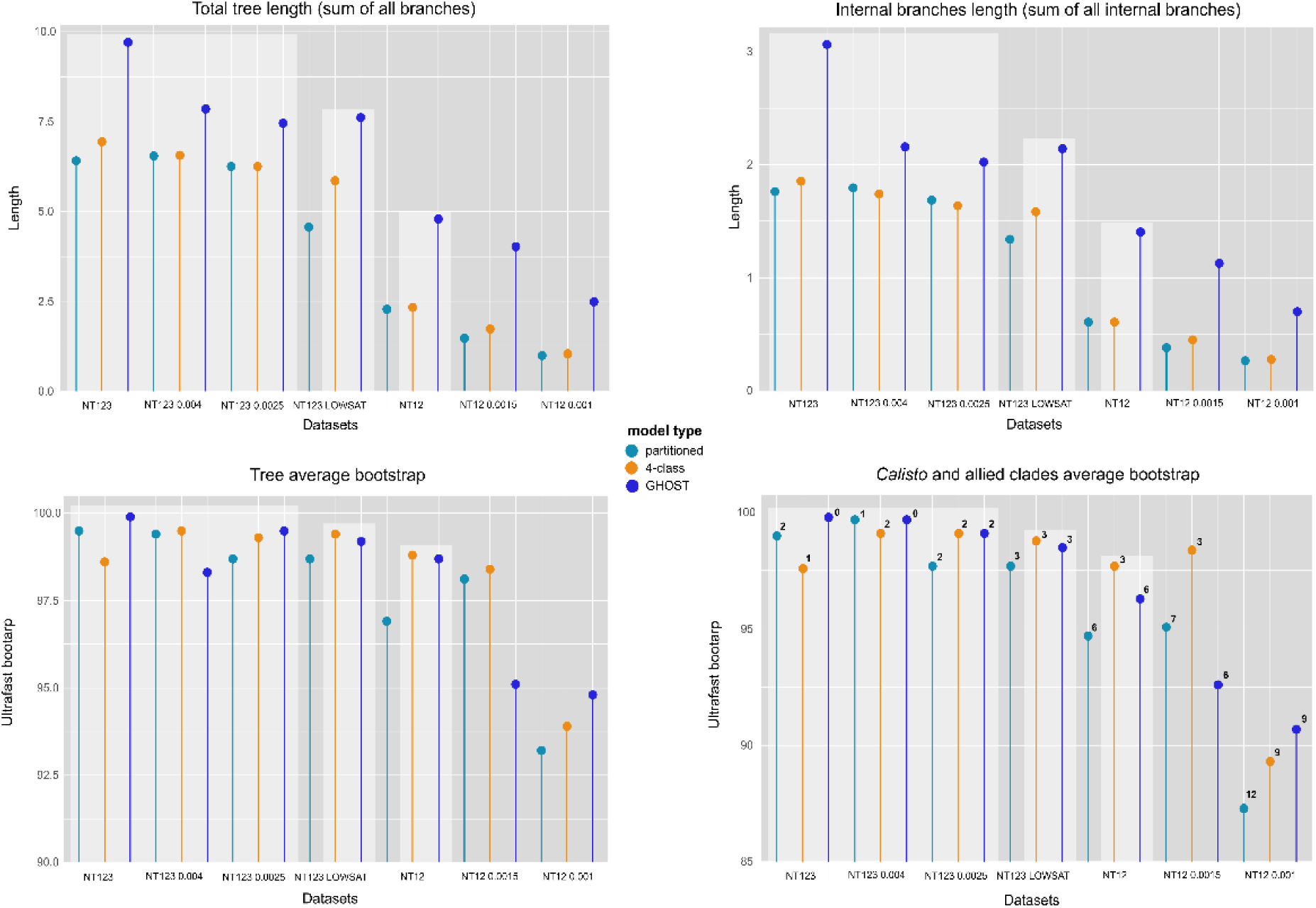
Comparison among trees inferred in IQ-TREE using all nucleotide datasets, except the amino acids ones, and partitioned and mixture models. Paler boxes mark trees with the same strongly supported topology. Numbers at the top of each lollipop in the bottom right plot represent the number of weakly supported branches. Data extracted from the IQ-TREE .iqtree files and the maximum likelihood tree.

We obtained relatively low mean values of both the gene concordance factor (gCF) and the site concordance factor (sCF) compared to the ultrafast IQ-TREE bootstrap values (File S4). The mean gCF and sCF values for the NT123 dataset were 61.6 and 52.5, respectively (Fig. S4, File S5). Concordance factors for the branches leading to splits between *Calisto* and its relatives were lower than these means, except for the Callerebiina–Ypthimina split, which had a sCF of 91.1.

### Incomplete Lineage Sorting and Ancestral Gene Flow

Aphid detected ancestral gene flow (GF) and incomplete lineage sorting (ILS) in the relationships inferred for *Calisto* and its closest relatives. ((*Calisto*,Erebiina)Pronophilina-Euptychiina)), with Erebiina also including Callerebiina, *Gyrocheilus*, Maniolina, andina, was the dominant topology in 615 of the 676 triplets containing 50 to 434 gene trees (Fig. S5). This topology is present in 44.6% of all gene trees in all triplets (File S6). When setting it as the main topology, the program detected an average 28.1% of no events across all assessed triplets and estimated 19.3% and 35.2% for ILS and GF conflict, respectively (Fig. S5). The second most common topology in the gene trees, ((*Calisto*, Pronophilina-Euptychiina) Erebiina), caused the highest posterior imbalance due to ILS and GF, with 59.6% and 66.9% respectively.

When the alternative topologies ((Erebiina, Pronophilina–Euptychiina) *Calisto*) and ((*Calisto*, Pronophilina–Euptychiina) Erebiina) were each specified as the main topology, aphid inferred average discordance due to ILS and GF of 19.7% and 59.1%, respectively, in the first case, and 21.6% and 45.8%, respectively, in the second. As expected, in both scenarios the topology most frequently recovered in the phylogenetic analyses, ((*Calisto*, Erebiina) Pronophilina–Euptychiina), and across most gene trees, explained between 58% and 99% of the discordance attributed to ILS and GF.

### Divergence Times and Ancestral Range Estimation

The three time-calibrated trees, inferred from different datasets and models and with different inner branch lengths, yielded relatively similar ages for the split between *Calisto* and its relatives, with most dates differing by 2-3 Myr and partially overlapping their 95% confidence intervals (Fig. 4, File S7). Our estimates suggest a late Oligocene - early Miocene split between the ancestors of *Calisto* and its close relatives.

**Figure 4.**
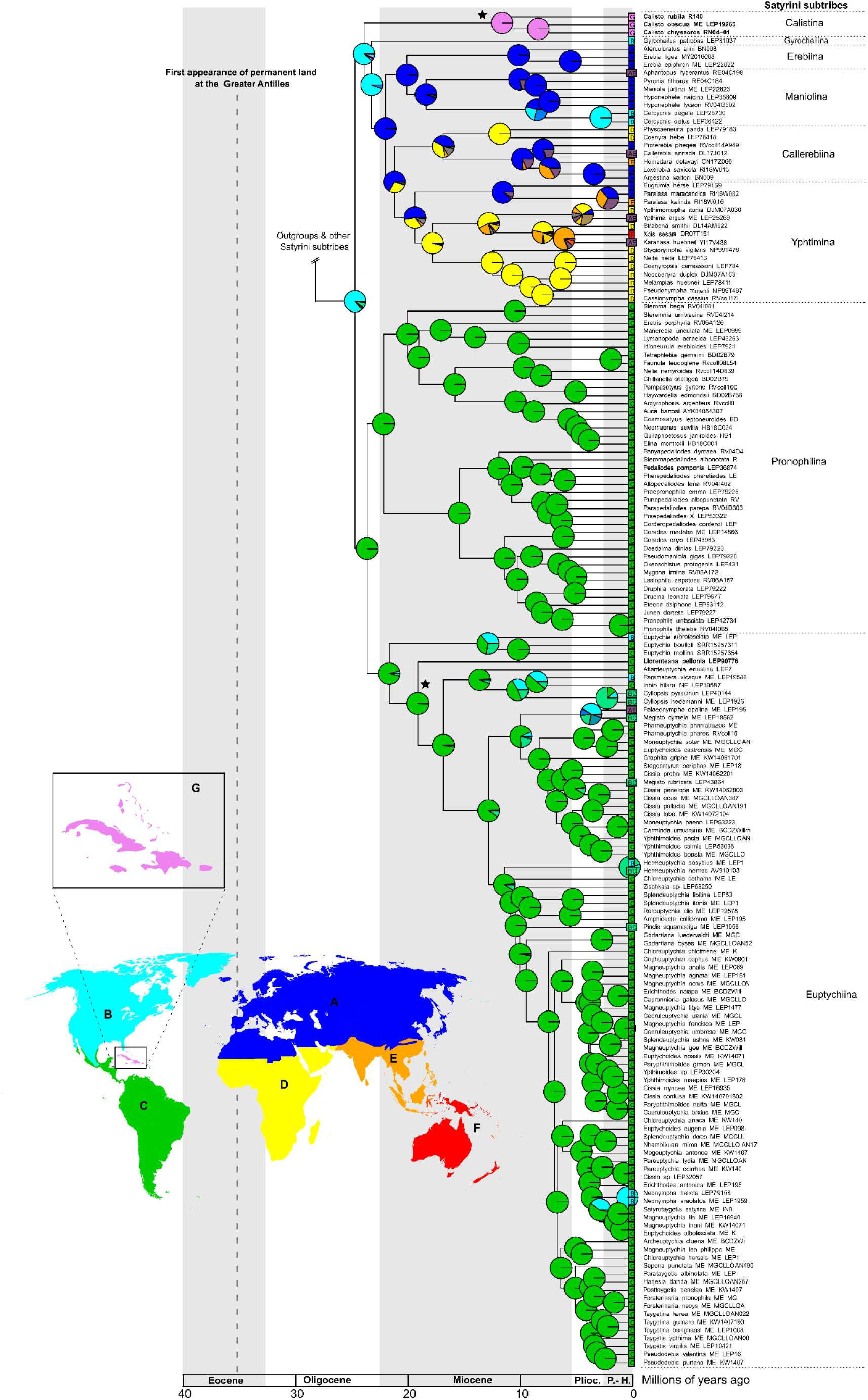
(Next page). BioGeoBEARS ancestral range reconstruction of the Satyrini subtribes closely related to *Calisto* and *Llorenteana* based on the maximum likelihood tree secondary inferred with the NT123 KW dataset and the GHOST model calibrated in RelTime with secondary calibration points from Espeland et al. (2018). The results of the best model (DEC + J) are represented. The black stars mark the positions of *Calisto* and *Llorenteana*. Areas: A, the Palearctic; B the Neartic; C, the Neotropics excluding the West Indies; D, the Afrotropics; E, the Indomalayan; F, Australasia; G, the West Indies, including the Greater Antilles, the Bahamas and Anegada Island (the entire range of *Calisto*). Refer to files S2 and S7 for the relative probabilities at nodes, the models comparison table, the species distribution per area, and the dispersal probabilities.

The constrained analysis with the DEC+J model had the highest relative probability, 99.9%, based on the AICc values and their weights (Fig. 4, File S7). This model estimated a Nearctic origin for the *Calisto* ancestor with 96.9% probability. Overall, the analysis suggests an Old World, likely Palearctic, origin for all clades sister to the subtribe Satyrina, whose ancestors colonized the New World from north to south. The analysis estimated with high probability the ancestral areas for *Gyrocheilus* (Nearctic), Erebiina (Palearctic), as well as for the two Neotropical subtribes, Pronophilina and Euptychiina, including *Llorenteana* (Neotropical) (Fig. 4, File S7). All models in the unconstrained analysis scored lower likelihood and AICc values recovering unlikely ancestral areas in some cases, *e.g.* a Palearctic origin for *Calisto* (File S7).

## Discussion

### *Mixture Models and the Phylogenetic Position of Calisto and* Llorenteana

Our study is the first to provide a comprehensive assessment of the phylogenetic position of the enigmatic genus *Calisto* and, for the first time, the placement of *Llorenteana* using molecular data. Previous studies on the Satyrini, as well as on most butterfly or Lepidoptera studies, have relied solely on traditional partitioned substitution models. Although alignment partitioning can improve the results of phylogenetic inference (McGuire et al. 2007; Tarasov and Dimitrov 2016), these models may violate pre-existing assumptions (Kolaczkowski and Thornton 2004; Ho 2009) or be too simplistic to recover all the evolutionary information of the sequences contained in a given alignment (Redmond and McLysaght 2021; Ren et al. 2025).

In most cases, our trees inferred in Astral-IV and IQ-TREE using partitioned models place *Calisto* in clade comprising the subtribes Callerebiina, Erebiina, Maniolina, *Gyrocheilus*, and Ypthimina. However, the relationships within this group inferred with partitioned models vary and lack support at several branches, with paraphyly appearing often. These results are similar to the obtained in previous studies, in which *Calisto*’s position changed depending on the dataset, sampling or the inference method used (*e.g.* Peña et al. 2011; Espeland et al. 2018), sometimes with the different placements receiving strong support (*e.g. Calisto* sister to *Gyrocheilus* and *Calisto* sister to *Euptychia* in Kawahara et al. 2023). Only the IQ-TREE trees inferred with the most informative datasets, NT123 and its versions with reduced compositional bias, fully resolved these relationships. We confirmed those relationships after applying 4-class mixture and GHOST models to these and the other nucleotide datasets. Only the least informative ones failed to recover them.

Overall, our results suggest that mixture models, both the 4-class and GHOST models, outperform the traditional partitioned models, echoing the results of prior intensive testing (Crotty et al. 2020; Ren et al. 2025). Most trees inferred with these models produced longer trees with strongly supported, longer inner branches and converged on a single topology, with the exception of the two least informative datasets. According to Ren et al. (2025), mixture models likely infer more of the substitutions underlying empirical alignments, resulting in trees of increased length and suggesting that partitioned models may fail to adequately account for the complexity of molecular evolution in phylogenomic studies. As per model fit, tests allowing an objective selection among partitioned and mixture models are not available yet (Crotty and Holland 2022; Liu et al. 2023). Despite limitations of its implementation for large genomic datasets, mixture models seem worth considering in the phylogenetic inference process alongside with the traditional partitioned models.

We found evidence of GF and ILS affecting our inference process, which could explain the differing hypotheses obtained in previous studies and in some of our results. Our analyses in aphid detected both phenomena independently of the main topology set. However, these analyses also showed that when the topology was set to our preferred relationships (the most common across our analyses) there was less conflict derived from ILS and GF than when setting the alternative topologies. These results could also partially explain the low gCF and sCF. Studies on other groups also affected by ILS and GF obtained similar results (Galtier 2024; Reboud et al. 2025).

Multispecies coalescent methods can handle missing data and polytomies in the input gene trees (Mirabab et al. 2021), but also assume that multiple sequence alignments and that gene trees are error free. Solís-Lemus et al. (2016) found that ASTRAL can be affected by anomalous gene trees due the presence of GF or a combination of ILS and GF. Our analyses in aphid showed that both phenomena are present in our datasets, and we believe that, together with other sources of gene tree error, such as model misspecification, they may have contributed to the inconsistencies observed in our ASTRAL trees.

### *The Origin of* Calisto *and* Llorenteana

Our ancestral area reconstruction analyses point toward a Nearctic origin for the ancestor of *Calisto* with the split from the ancestor of continental relatives likely occurring around 24 Ma. These results support the hypothesis proposed by Peña et al. (2011), that Satyrini ancestors colonized the New World from north to south. All members of *Calisto*’s sister clade inhabit the Nearctic and the Old World, and while the branch leading to Pronophilina and Euptychiina has a Neotropical origin, it is preceded by a branch suggesting an ancestor of Nearctic origin. The latter is supported by the Nearctic or extreme northern Neotropical distributions of *Llorenteana* and some *Euptychia*, which represent the earliest splits within Euptychiina (Viloria and Martinez 2019; Espeland et al. 2023).

These results conflict with the hypothesis by Matos-Maravi et al. (2014) of a colonization event from South America via GAARlandia. This hypothesis proposes that a landspan, a chain of islands, or a mix of both, interconnected South America with the islands serving as colonization route for ancestors of the current biota of the islands until *ca.* 35-33 Ma (Iturralde-Vinent and MacPhee 1999; Iturralde-Vinent 2006). This theory has been used to explain the arrival of several groups in the Caribbean (Iturralde-Vinent 2006; Alonso et al. 2012; McHugh et al. 2014; Tong et al. 2018). However, other studies have reached different conclusions (Sato et al. 2016; Price et al. 2022) or questioned its validity altogether (Ali 2012; Ali and Hedges, 2024; Massip-Veloso et al. 2024). Although the inferred geographical area of origin of *Calisto*’s ancestor is not close to South America, other possible pathways may have existed. Available geological reconstructions include landmasses and shallow platforms extending from Central America toward the Caribbean Sea and the archipelago since the late Eocene, such as the Mayan Block and the Pedro Bank (Iturralde-Vinent 2006). Although these structures apparently never formed a direct connection, they may have facilitated the movement of the biota by reducing the distance between the islands and the continent.

### On the Subtribal Classification of Calisto and Allied Groups

Zhang et al. (2021) established several new subtribes within Satyrini to accommodate orphaned taxa. They named Callerebiina to include several genera formerly placed in Ypthimina, and Calistina and Gyrocheilina to accommodate each a single genus, *Calisto* and *Gyrocheilus* respectively. The authors justified the subtribal status of *Calisto* based on its position as sister to Euptychiina, albeit weakly supported, as well as on wing venation characters. However, all mentioned diagnostic characters occur in other Satyrini. For instance, the forewing radial veins also arise in a single branch from the cell in *Argyrophorus*, *Argyrophenga*, *Llorenteana*, *Megisto* and in some species of *Yphtima* (Miller 1968; Viloria 2007). For *Gyrocheilus*, the authors mention a full generic diagnosis, whereas for the clade containing *Callerebia* and related genera, they relied only on molecular characters. Consequently, the parent clade containing all these taxa now comprises six subtribes: Calistina, Gyrocheilina, Callerebiina, Erebiina, Maniolina, and Ypthimina.

In our opinion, this degree of splitting within a natural clade is unnecessary and introduces instability to the already complex Satyrini nomenclature. This approach also contrasts with the status of the two clades sister to the aforementioned one, each of which contains a single subtribe: Pronophilina and Eutptychiina. Considering the strongly supported relationships recovered and the biogeographical results, it would be more consistent to recognize a single subtribe of predominantly Old Word distribution that includes a few genera that colonized or evolved in the Nearctic, such as *Cercyonis*, *Erebia*, and *Gyrocheilus*, as well as *Calisto* in the West Indies. Similarly, both Euptychiina and Pronophilina are predominantly Neotropical, with a few taxa such as *Euptychia*, *Cyllopsis*, *Megisto* and *Paramacera* reaching the Nearctic, and *Megisto* furthermore dispersing from the Nearctic to the Old world. Therefore, we propose recognizing a single subtribe for *Calisto* and all allied taxa. Because the name Erebiina (Tutt 1896) predates Maniolina and Ypthimina (both Reuter 1897), it has priority, and all other proposed subtribal names for this clade should be considered its junior synonyms: Calistina **syn. nov.**, Callerebiina **syn. nov.**, Maniolina **syn. nov.**, and Ypthimina **syn. nov.** This clade, like many others within Satyrinae, currently lacks a clear morphological diagnosis and will remain so pending further integrative analyses combining well-resolved phylogenies with morphological data.

## Funding Acknowledgement

This work was funded by the Deutsche Forschungsgemeinschaft (DFG) NU 452/1-1.

## Acknowledgements

We are grateful to Lars Podsiadlowski and Sandra Kukowka for their help in the molecular lab at the LIB, Bonn. We thank Keith R. Willmott, Andrew Warren, Deborah Matthews Lott and Andrei Sourakov for their help during the visit to the McGuire Center for Lepidoptera. RN acknowledges funding by the Deutsche Forschungsgemeinschaft (DFG) NU 452/1-1.

## Data Availability Statement

We deposited on GenBank all new genomic sequencing raw data produced during this study (PRJNA1402963). All datasets, tree files and data used for analyses can be found on Dryad, DOI:10.5061/dryad.31zcrjf25, and Zenodo, 10.5281/zenodo.18234177.

## Conflict of Interest Statement

None declared.

---

**Table S1.**
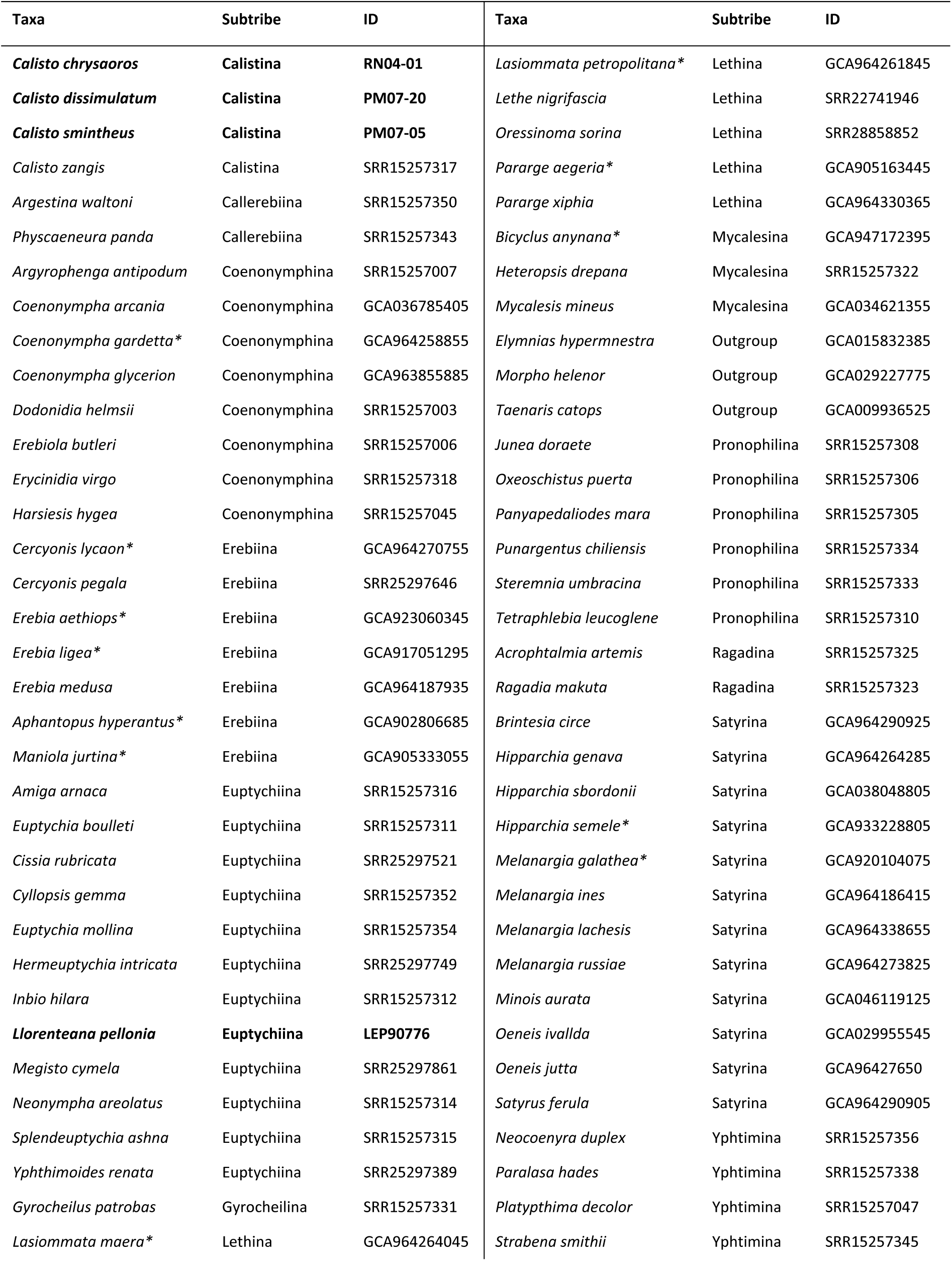
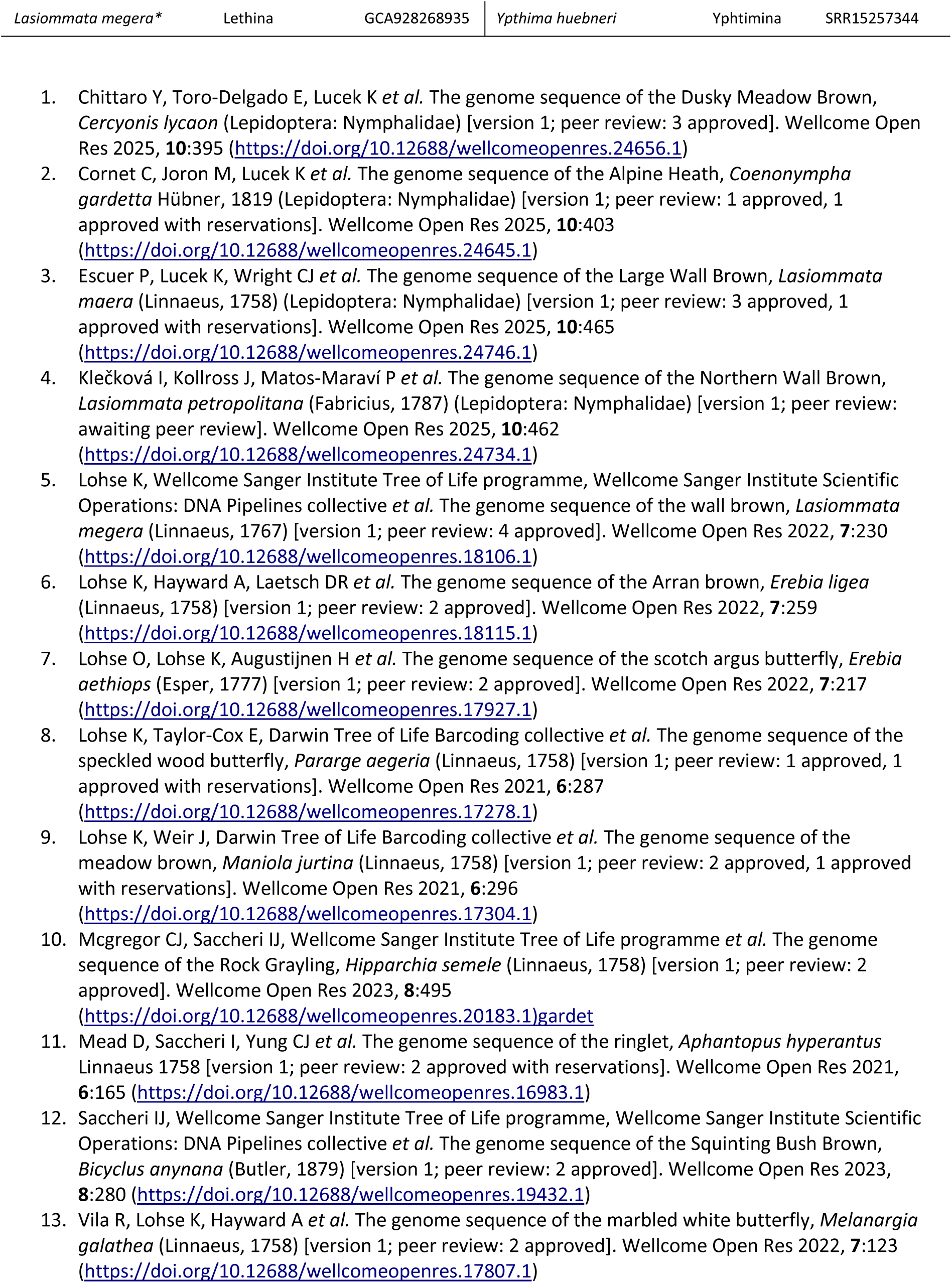
Specimens included in this study. Newly produced samples shown in bold. * Genome assemblies generated by the Darwin Tree of Life project (The Darwin Tree of Life Project Consortium 2022) and Project Psyche (Wright et al. 2025) whose genome notes appear listed below.

**Table S2.**
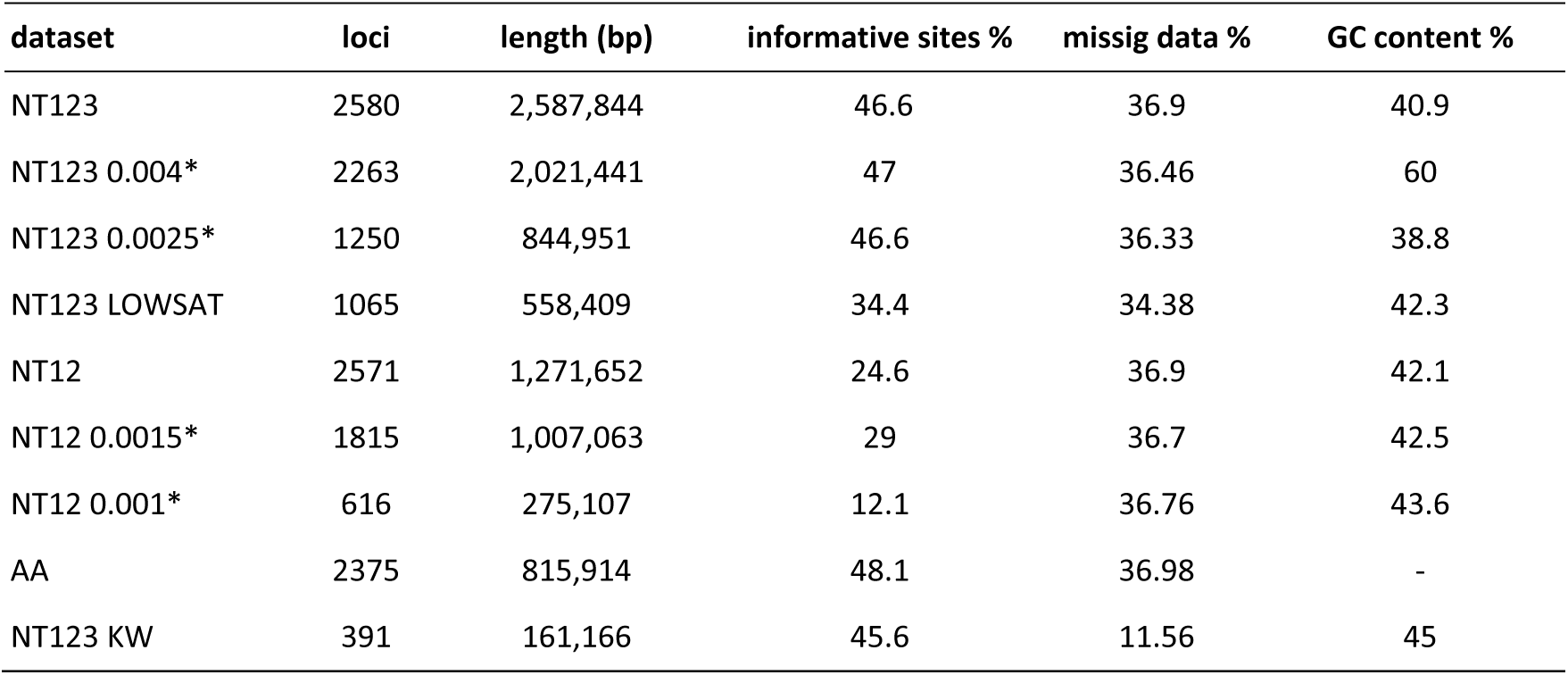
Datasets statistics. *normalized Relative Compositional Frequency Variation (nRCFV) threshold value used to discard genes deviating from compositional homogenenity (see main text)

**Figure S1.**
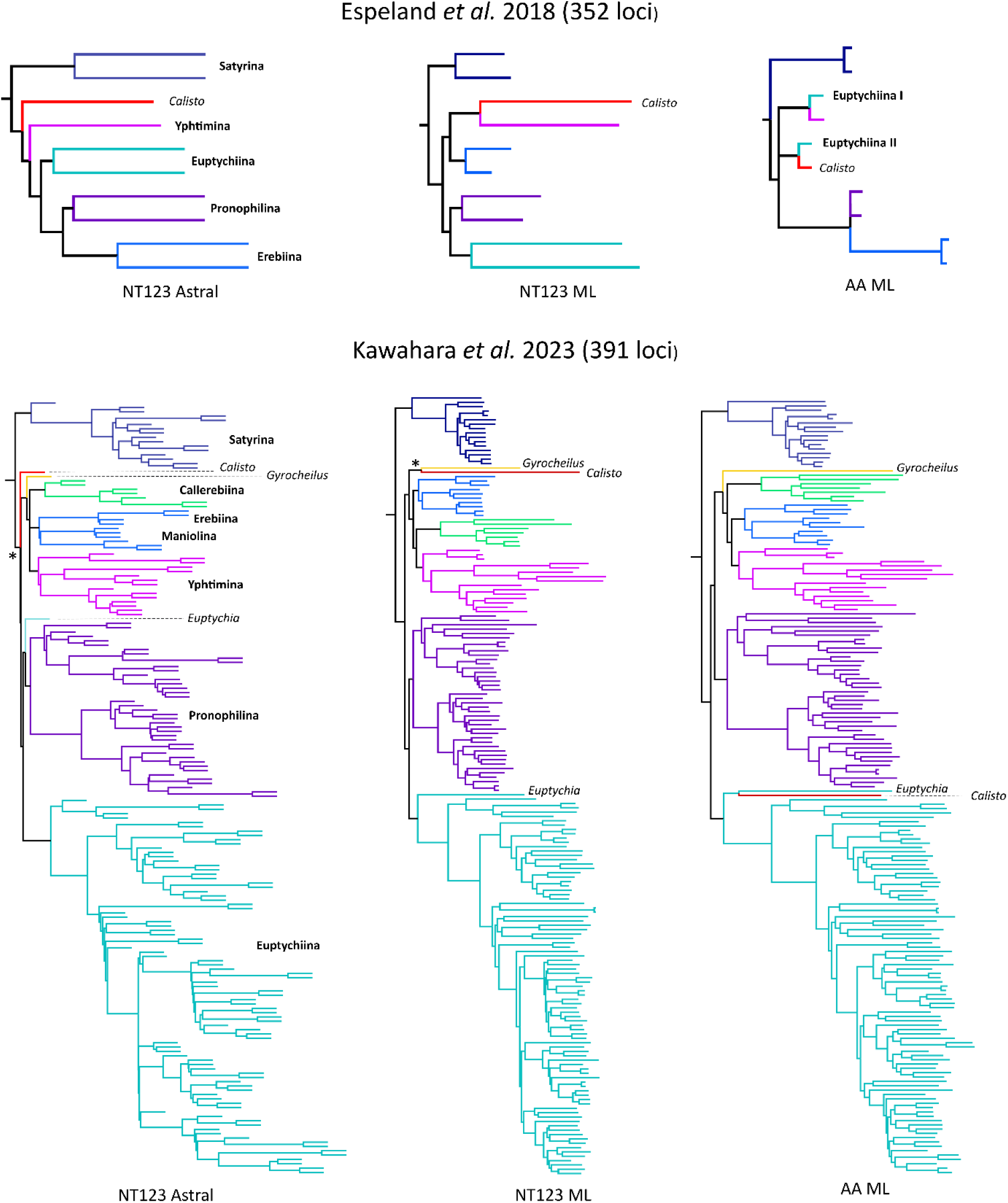
Examples of different phylogenetic placements inferred for *Calisto* by Espeland et al. (2018) and Kawahara et al. (2023). Colors and names identify closely related subtribes and the genera *Calisto*, *Euptychia*, and *Gyrocheilus*. Abbreviations and symbols: NT123- nucleotide, all codon positions, AA- amino acids, ML-maximum likelihood, *- strong support.

**Figure S2.**
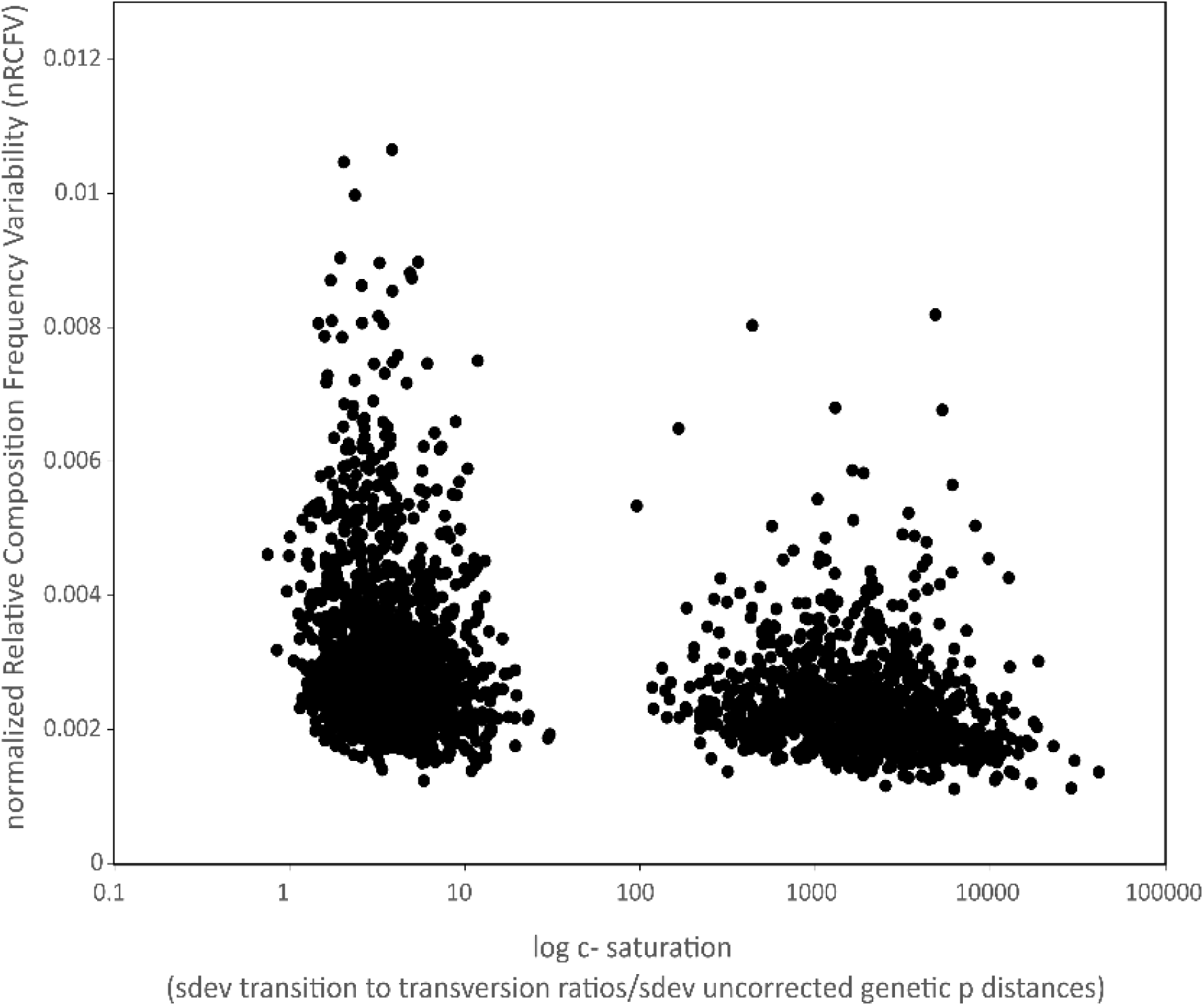
Normalized Relative Compositional Frequency Variation (nRCFV) and c- convergence saturation values calculated by BaCoCa for the 2580 genes included in the main dataset, NT123.

**Figure S3.**
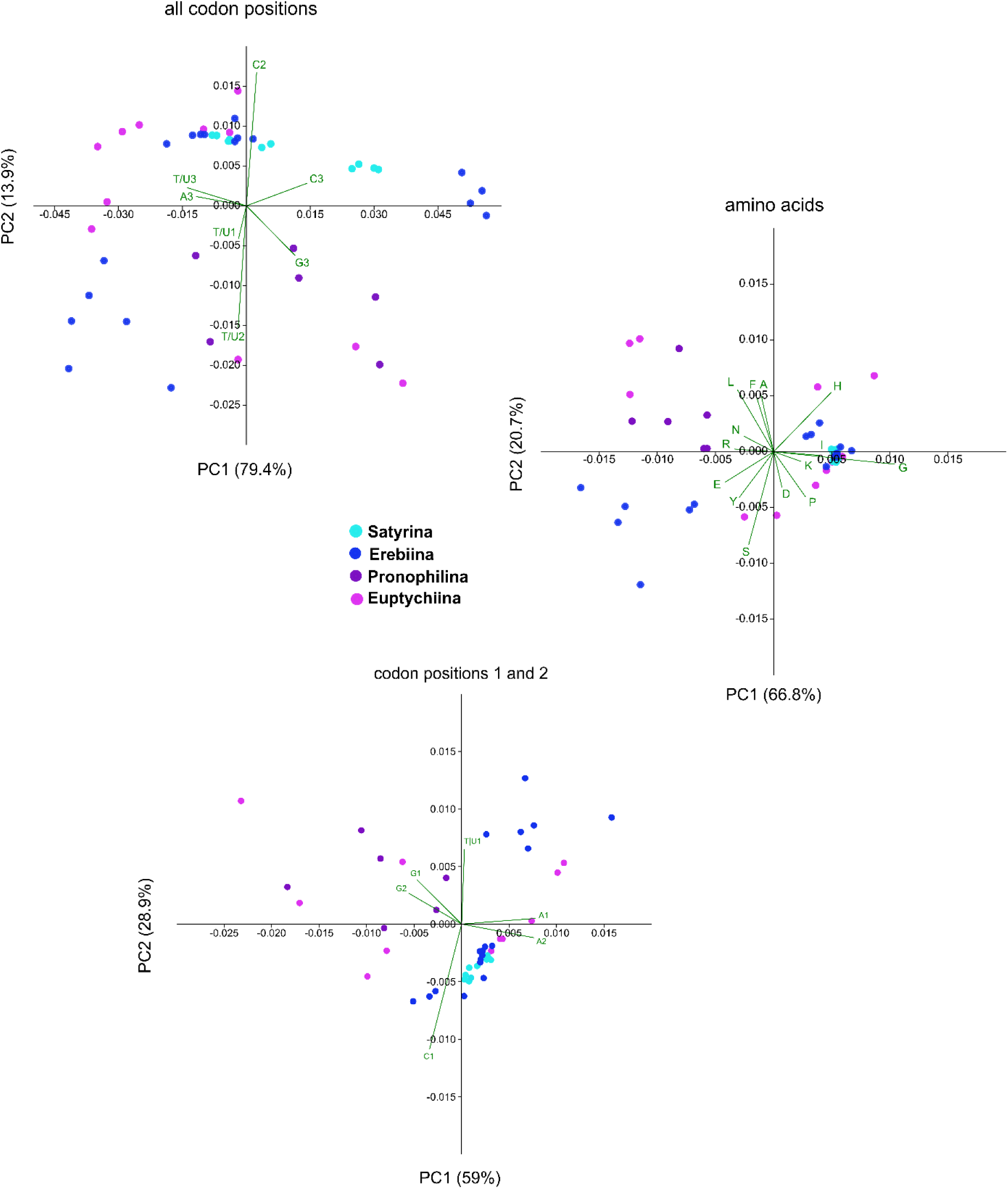
Principal Component Analyses of the base frequencies estimated in BaCoCa for the samples of *Calisto*, *Llorenteana*, and their closely related taxa. Colors identify *Calisto* related subtribes as shown, for practical reasons Erebiina also includes Calistina, Callerebiina, Gyrocheilina, Maniolina, and Ypthimina. The biplot at each plot shows the codon positions and amino acids contributing the most to the variability observed.

**Figure S4.**
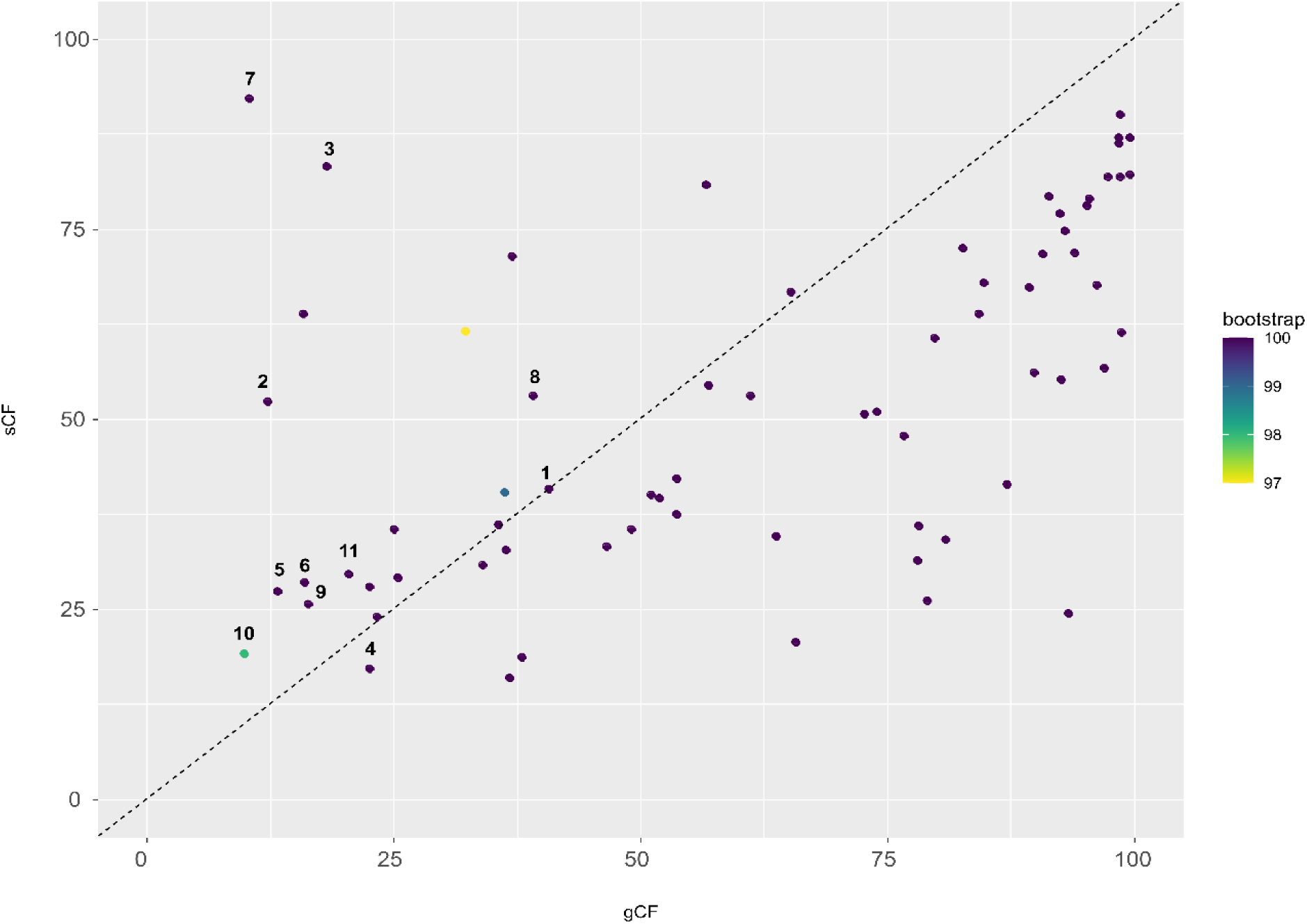
Gene (gCF) and site concordance factor (sCF) values calculated in IQ-TREE using the NT123 tree inferred with the GHOST model. Each point represents a node of the phylogeny and its color represents the ultrafast bootstrap value (bootstrap). The dashed line corresponds to the identity line. Relevant nodes: 1 (Pronophilina, Euptychiina), (*Calisto* (*Gyrocheilus* (Erebiina-Maniolina (Callerebiina, Ypthimina)))), 2 (Pronophilina, Euptychiina), 3 Pronophilina, 4 Euptychiina, 5 (*Calisto* (*Gyrocheilus* (Erebiina-Maniolina (Callerebiina, Ypthimina)))), 6 (*Gyrocheilus* (Erebiina-Maniolina (Callerebiina, Ypthimina))), 7 (Callerebiina, Ypthimina), 8 Callerebiina, 9 Ypthimina, 10 (Erebiina-Maniolina (Callerebiina, Ypthimina)), 11 Erebiina-Maniolina.

**Figure S5.**
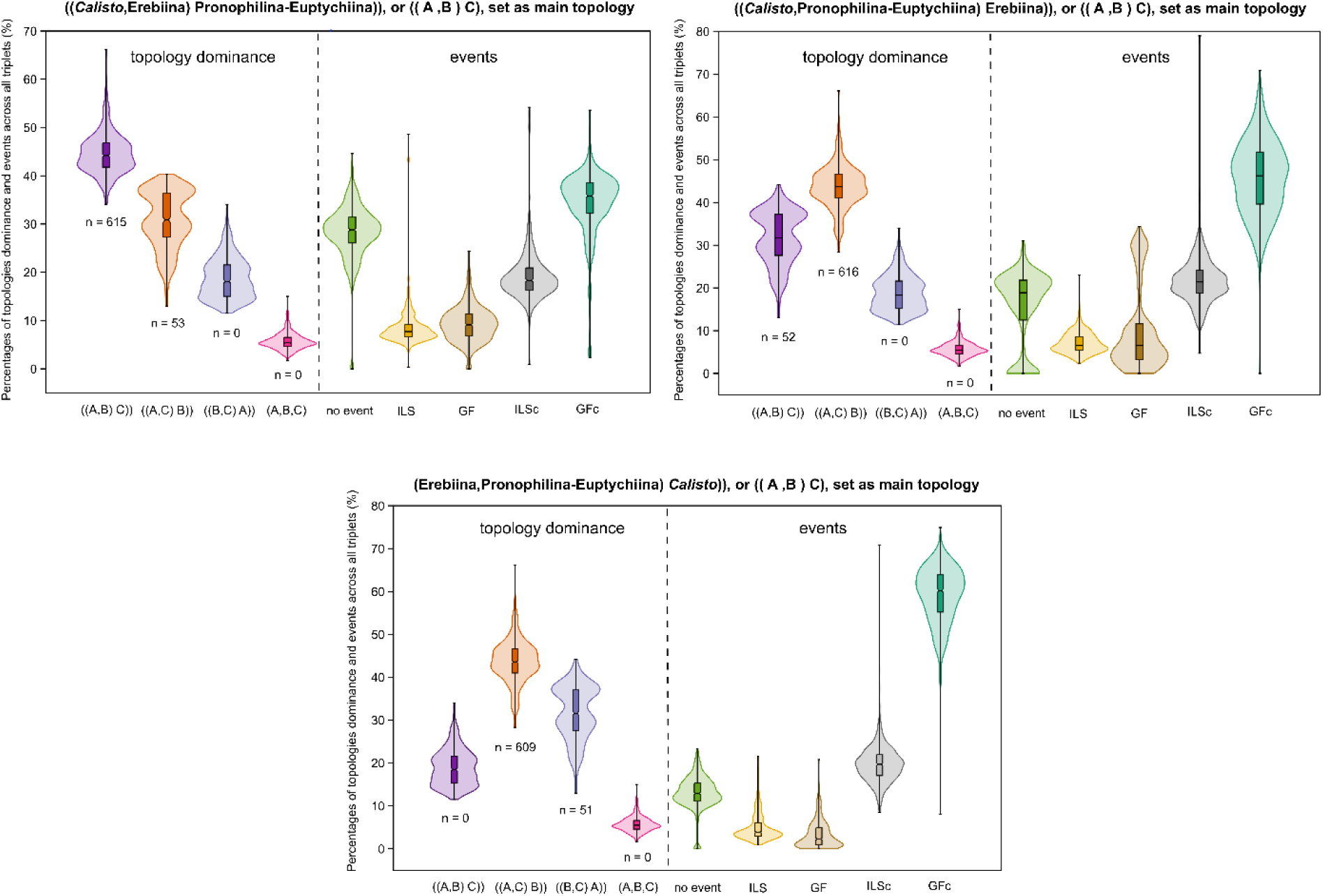
Summary of all aphid analyses. Percentage of the dominance of each topology across all triplets analyzed when setting three alternative topologies and the percentage of no events detected (no event), incomplete lineage sorting events (ILS), gene flow events (GF), incomplete lineage sorting events causing posterior conflict (ILSc), and gene flow events causing posterior conflict (GFc). The “n” below each violin represents the number of triplets in which each topology was the dominant. Erebiina also includes Callerebiina, Gyrocheilina, Maniolina, and Ypthimina.

## Notes

https://zenodo.org/records/18234177?token=eyJhbGciOiJIUzUxMiJ9.eyJpZCI6Ijc0NTcyYTcxLTBhNjQtNGVhMS04YzE1LWVkZDYwNDdlMGMyNiIsImRhdGEiOnt9LCJyYW5kb20iOiJiZDc2OWJjMmQxY2JmNTFjNzFlZTNhOTgyM2ZlYTk3MCJ9.Ox1_rOXgVx4Wia4e_vipiCDisp-JpUYJFsRPimspYyKeeixGo9R7C28tQOK1OwiYHoNiOV8nQjteD-EFVa-3CQ

### Competing Interest Statement

The authors have declared no competing interest.

